# Intratumoral heterogeneity in microsatellite instability at single cell resolution

**DOI:** 10.1101/2025.06.11.658021

**Authors:** Harrison Anthony, Cathal Seoighe

## Abstract

Subclonal diversity within a tumor is highly relevant for tumor evolution and treatment. This diversity is often referred to as intratumoral heterogeneity and is known to complicate the interpretation of single-test biomarkers. Microsatellite instability (MSI) is one such biomarker, which is used to guide immune check-point inhibitor treatment through the classification of samples as either having high microsatellite instability (MSI-H) or as being microsatellite stable (MSS). Although established as a therapeutic biomarker, it remains unclear whether MSI itself is a heterogeneous phenomenon. To investigate heterogeneity in MSI status, we integrated single-cell sequencing data from 134 samples across 49 individuals and developed a computational pipeline to infer MSI-H cells and quantify heterogeneity in MSI status. We found evidence of intratumoral heterogeneity in MSI both in individuals originally classified as MSI-H and MSS. Approximately a third of individuals showed evidence of divergence in MSI status between distinct clusters of cancer cells and most individuals had distinct MSI-H and MSS subclones. These results challenge the assumption that MSI should be treated as a binary biomarker and suggest the single-biopsy tests in current use could overlook a salient feature of this important molecular phenotype. Accounting for heterogeneity may lead to improved biomarker performance and, potentially, help explain reports of intrinsic treatment resistance and low overall response rate in MSI-H cancers. Further studies are warranted to determine the frequency of heterogeneity in MSI at the population level, and whether the presence of both MSI-H and MSS subclones can have clinical implications.

## 1 Background

Subclonal diversity within a tumor is a critical consideration in cancer research and treatment. The overall diversity found in a single neoplasia is called intratumoral heterogeneity (ITH). While ITH was first conceptualized to be genetic in nature [1], it is now used to describe genetic, epigenetic, and phenotypic differences between sub-clones [2]. The diversity within a tumor is important because ITH has been linked to poor patient outcomes, therapy resistance, and relapse [3, 4]. Furthermore, biomarkers that rely on single-sample tests can be susceptible to sampling bias when ITH is present [5, 6]. While its origins are still debated [7], one well known driver of ITH is genomic instability [8].

Genomic instability is a hallmark of cancer, characterized by a higher rate of accumulation of mutations during replication, typically due to deficiencies in DNA repair genes [9]. The two most common forms of genomic instability are at the chromosomal level, where instability is characterized by aneuploidy and chromosomal aberrations [10], and at the microsatellite level, where short tandem repeats expand and contract in a mutator phenotype manner [11]. The latter, referred to as microsatellite instability (MSI), is hypothesized to be the result of a deficient mismatch repair (dMMR) pathway and is commonly used as a biomarker to help guide immune checkpoint inhibitor treatment. This is done by classifying cancers as either having high microsatellite instability (MSI-H) or as being microsatellite stable (MSS) [12]. The classification is normally carried out using a single-sample test that compares five microsatellite markers between a tumor and paired-normal sample [13]. While the interplay between chromosomal instability and ITH is well defined and explained [14–16], the relationship between MSI and ITH is less clear.

Up to this point, most research on MSI and ITH has been framed around how MSI can impact and shape the variation present within a tumor. Most notable have been studies focusing on specific mutations [17, 18] and the immune cell types present in the tumor microenvironment [19, 20]. Although, there have been reported cases of ITH in MSI status [21–26], the question of whether MSI itself is frequently a heterogeneous phenomenon, with some subclones displaying MSI while others do not, has yet to be examined in detail. This warrants further investigation as it may help to explain limitations of MSI-H as a biomarker for precision medicine, such as low treatment response rates and intrinsic treatment resistance [27–29]. Taking heterogeneity into account may, ultimately, lead to improved biomarker performance.

The current literature suggests that subclonality of MSI status is relatively rare or entirely absent [30, 31], but that is not always the case. There are many examples of individuals not only with discordant MSI statuses between the primary tumor site and metastases [21–24] but also between multiple sites in the primary tumor [25, 26]. While these are small case studies, they provide anecdotal evidence for cancers comprising MSI-H and MSS subclones. However, there has, as yet, been no attempt to evaluate the frequency with which this occurs. Furthermore, a detailed examination of heterogeneity requires an assessment of MSI at the single-cell level with next-generation sequencing, not with the traditional methods of PCR and IHC used in these case studies, as these are limited to detecting clear spatial heterogeneity. Here we aimed to address these gaps through an analysis of published single-cell datasets that include paired clinical MSI status. To do this, we developed a custom Snakemake [32] pipeline that identifies MSI-H cells and uses novel methods to assess levels of heterogeneity and have made this pipeline available as an open-source, scalable resource to the scientific community. We evaluated the pipeline by mixing varying numbers of MSI-H and MSS cells from different samples. Applying this framework, we show evidence of heterogeneity in MSI status at the single-cell level and estimate its prevalence in the curated data. We also examine the nature of MSI heterogeneity through a detailed investigation of single-cell data from two individuals – one classified as MSI-H and the other MSS through PCR/IHC tests.

## 2 Methods

### 2.1 Datasets

We used single-cell RNA sequencing data that was generated as part of three previous studies [33–35] (Table 1). Raw FASTQ files were downloaded from either the European Genome-Phenome Archive (RRID:SCR004944) or from the Sequence Read Archive (RRID:SCR001370). All other data was downloaded in matrix format from the Gene Expression Omnibus (RRID:SCR005012). The data consists of 134 samples from 49 individuals with metastatic or non-metastatic colorectal cancer. Individuals were grouped into MSI-H and MSS categories based on the original PCR/IHC clinical status reported in previous studies. In total there were 29 deemed MSI-H, 18 MSS, and two did not have a reported MSI status (Table 2). Each sample was created with either Single Cell 3’ v2, 3’ v3, or 5’ Reagent Kit from 10X Genomics and was sequenced either on an Illumina NextSeq 500, NovaSeq 6000, BGISEQ DNBSEQ-T7, or HiSeq X Ten machine. Complete sequencing and library preparation information can be found by referencing the Dataset ID in (Table 1). Individuals from datasets EGAD00001008555, EGAD00001008584, EGAD00001008585 had multi-regional samples from the same tumor and multi-site samples from metastatic tissue and lymph nodes. Even though variation in the multi-site samples could be considered intra-individual heterogeneity rather than ITH, we kept them in the analysis to retain as many cancer cells and as much heterogeneity as possible. The other two datasets GSE205506 and PRJNA932556 include some individuals that had treatment for MSI (anti-PD-1 and celecoxib). We excluded the following samples because we did not identify any cancer cells: XHC080-SI-GA-B11, XHC082-SI-GA-C1, XHC127-SI-GA-F10, EXT129, EXT051, and EXT097.

**Table 1.** Individual summary statistics and subclone information.

| Dataset ID | Cancer type | Individuals | Samples | Sequencing |
| --- | --- | --- | --- | --- |
| EGAD00001008555 | Colorectal/metastatic | 15 | 77 | Illumina HiSeq 4000 |
| EGAD00001008584 | Colorectal/metastatic | 3 | 6 | Illumina HiSeq 4000 |
| EGAD00001008585 | Colorectal/metastatic | 6 | 18 | Illumina NextSeq 500/NovaSeq6000 |
| GSE205506 | Colorectal | 19 | 27 | Illumina NovaSeq 6000/DNBSEQ-T |
| PRJNA932556 | Colorectal | 6 | 6 | HiSeq X Ten |

**Table 2.**
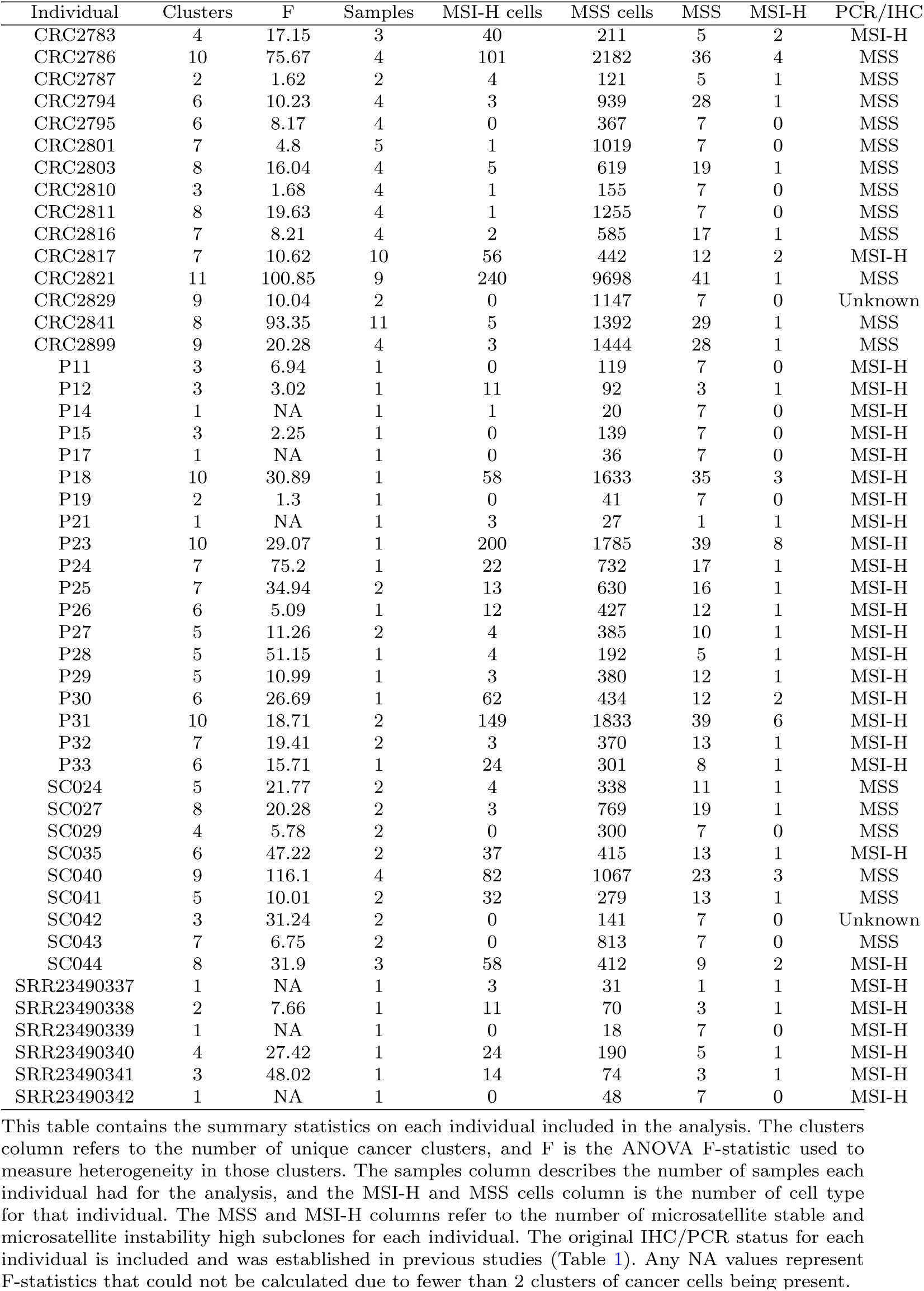
Individual summary statistics and subclone information.

| Individual | Clusters | F | Samples | MSI-H cells | MSS cells | MSS | MSI-H | PCR/IHC |
| --- | --- | --- | --- | --- | --- | --- | --- | --- |
| CRC2783 | 4 | 17.15 | 3 | 40 | 211 | 5 | 2 | MSI-H |
| CRC2786 | 10 | 75.67 | 4 | 101 | 2182 | 36 | 4 | MSS |
| CRC2787 | 2 | 1.62 | 2 | 4 | 121 | 5 | 1 | MSS |
| CRC2794 | 6 | 10.23 | 4 | 3 | 939 | 28 | 1 | MSS |
| CRC2795 | 6 | 8.17 | 4 | 0 | 367 | 7 | 0 | MSS |
| CRC2801 | 7 | 4.8 | 5 | 1 | 1019 | 7 | 0 | MSS |
| CRC2803 | 8 | 16.04 | 4 | 5 | 619 | 19 | 1 | MSS |
| CRC2810 | 3 | 1.68 | 4 | 1 | 155 | 7 | 0 | MSS |
| CRC2811 | 8 | 19.63 | 4 | 1 | 1255 | 7 | 0 | MSS |
| CRC2816 | 7 | 8.21 | 4 | 2 | 585 | 17 | 1 | MSS |
| CRC2817 | 7 | 10.62 | 10 | 56 | 442 | 12 | 2 | MSI-H |
| CRC2821 | 11 | 100.85 | 9 | 240 | 9698 | 41 | 1 | MSS |
| CRC2829 | 9 | 10.04 | 2 | 0 | 1147 | 7 | 0 | Unknown |
| CRC2841 | 8 | 93.35 | 11 | 5 | 1392 | 29 | 1 | MSS |
| CRC2899 | 9 | 20.28 | 4 | 3 | 1444 | 28 | 1 | MSS |
| P11 | 3 | 6.94 | 1 | 0 | 119 | 7 | 0 | MSI-H |
| P12 | 3 | 3.02 | 1 | 11 | 92 | 3 | 1 | MSI-H |
| P14 | 1 | NA | 1 | 1 | 20 | 7 | 0 | MSI-H |
| P15 | 3 | 2.25 | 1 | 0 | 139 | 7 | 0 | MSI-H |
| P17 | 1 | NA | 1 | 0 | 36 | 7 | 0 | MSI-H |
| P18 | 10 | 30.89 | 1 | 58 | 1633 | 35 | 3 | MSI-H |
| P19 | 2 | 1.3 | 1 | 0 | 41 | 7 | 0 | MSI-H |
| P21 | 1 | NA | 1 | 3 | 27 | 1 | 1 | MSI-H |
| P23 | 10 | 29.07 | 1 | 200 | 1785 | 39 | 8 | MSI-H |
| P24 | 7 | 75.2 | 1 | 22 | 732 | 17 | 1 | MSI-H |
| P25 | 7 | 34.94 | 2 | 13 | 630 | 16 | 1 | MSI-H |
| P26 | 6 | 5.09 | 1 | 12 | 427 | 12 | 1 | MSI-H |
| P27 | 5 | 11.26 | 2 | 4 | 385 | 10 | 1 | MSI-H |
| P28 | 5 | 51.15 | 1 | 4 | 192 | 5 | 1 | MSI-H |
| P29 | 5 | 10.99 | 1 | 3 | 380 | 12 | 1 | MSI-H |
| P30 | 6 | 26.69 | 1 | 62 | 434 | 12 | 2 | MSI-H |
| P31 | 10 | 18.71 | 2 | 149 | 1833 | 39 | 6 | MSI-H |
| P32 | 7 | 19.41 | 2 | 3 | 370 | 13 | 1 | MSI-H |
| P33 | 6 | 15.71 | 1 | 24 | 301 | 8 | 1 | MSI-H |
| SC024 | 5 | 21.77 | 2 | 4 | 338 | 11 | 1 | MSS |
| SC027 | 8 | 20.28 | 2 | 3 | 769 | 19 | 1 | MSS |
| SC029 | 4 | 5.78 | 2 | 0 | 300 | 7 | 0 | MSS |
| SC035 | 6 | 47.22 | 2 | 37 | 415 | 13 | 1 | MSI-H |
| SC040 | 9 | 116.1 | 4 | 82 | 1067 | 23 | 3 | MSS |
| SC041 | 5 | 10.01 | 2 | 32 | 279 | 13 | 1 | MSS |
| SC042 | 3 | 31.24 | 2 | 0 | 141 | 7 | 0 | Unknown |
| SC043 | 7 | 6.75 | 2 | 0 | 813 | 7 | 0 | MSS |
| SC044 | 8 | 31.9 | 3 | 58 | 412 | 9 | 2 | MSI-H |
| SRR23490337 | 1 | NA | 1 | 3 | 31 | 1 | 1 | MSI-H |
| SRR23490338 | 2 | 7.66 | 1 | 11 | 70 | 3 | 1 | MSI-H |
| SRR23490339 | 1 | NA | 1 | 0 | 18 | 7 | 0 | MSI-H |
| SRR23490340 | 4 | 27.42 | 1 | 24 | 190 | 5 | 1 | MSI-H |
| SRR23490341 | 3 | 48.02 | 1 | 14 | 74 | 3 | 1 | MSI-H |
| SRR23490342 | 1 | NA | 1 | 0 | 48 | 7 | 0 | MSI-H |

### 2.2 Data processing

We aligned FASTQ files to the GRCh38 human reference genome and converted them to a gene count matrix using the 10x Genomics Cell Ranger v7.2.0 software suite. From there all matrix files were processed following the Seurat best practices tutorials (https://satijalab.org/seurat/). Briefly, Seurat objects were created and only genes detected in a minimum of 3 cells were used for downstream analysis. We further filtered out cells with fewer than 100 features, more than 3000 features, and if a cell had more than 35 percent of all genes labeled as mitochondrial. While we followed the Seurat best practices closely, the default settings aim to maximize immune cell type identification and filtered out the majority of cancer cells. The filter settings described here were designed to maximize the number of tumor cells retained while still removing cells that were poor-quality or likely necrotic. These filter settings, specifically mitochondrial gene percentage, are supported by a recent study that showed that filter settings that are too strict remove viable cancer cells from single-cell sequencing data [36]. After filtering, all gene count matrices then were normalized using the LogNormalize option with the scale setting set to 10,000. The 2,000 most variable genes were found using the “vst” selection method and were used to cluster together groups of cells with the RunPCA function. The first 15 PCA dimensions were used to run the following functions: FindNeighbors, FindClusters (resolution set to 0.5), and RunUMAP. If an individual had multiple samples, they were integrated together by using the IntegrateLayers function, with the method set to CCAIntegration and k.weight set to 50. Each integrated sample then underwent re-clustering with the set-tings previously mentioned. The integration step after subsetting down to only cancer cells used the same settings except the FindClusters resolution was set to 0.8, and only the first 10 principal components were used.

### 2.3 Cell classification and measuring ITH

After each sample is processed, all cells are classified as either cancer or normal and then MSI-H or MSS. These classification steps are built upon two machine learning based programs trained on large pan-cancer datasets. The first, scATOMIC [37], was used to distinguish tumor cells from normal ones, and the second, MSIsensor-RNA [38] determined MSI status. Both tools were run with default settings, but to get an MSI score for each cell, we had to transform the prebuilt MSIsensor-RNA baseline file. This was done by filtering both the count matrix and baseline to only include gene names common to both. The filtered baseline and count matrix files were then used to get MSI scores for all cells within a sample. From there cells were classified as MSI-H if they also were labeled as cancer by scATOMIC and if they had an MSI score of .75 or more (75% probability the cell is MSI-H). Levels of ITH were assessed with two methods. First, we measured ITH by testing for differences in mean MSI score between cancer cell clusters with a one-way ANOVA test. The ANOVA F-statistic was used to describe levels of heterogeneity in the biomarker, with a large value of the F-statistic indicating greater heterogeneity in MSI. Secondly, we identified subclones within each individual by comparing CNVs between MSI-H, MSS, and normal cells. This was done by passing the relevant cell classification for each unique barcode to InferCNV [39] (DOI: 10.18129/B9.bioc.infercnv). InferCNV was used with default settings except in the case of CRC2821, which had many more cancer cells than the other samples. We increased the k nn setting from the default of 20 to 50 to take into account the larger dataset. Lastly, we ran differential expression using a Wilcoxon Rank Sum Test (the default for Seurat) between clusters of cancer cells and between MSI-H and MSS cancer cells for each individual. We verified how well our pipeline captured heterogeneity in MSI status by mixing together randomly sampled tumor and normal cells in varying proportions using a custom R script. We simulated varying levels of heterogeneity by mixing the cells of one sample that had homogenous MSI-H cancer cells (GSM6213995 from individual P33) and one sample with homogenous MSS cancer cells (XHC118-SI-GA-F1 from individual CRC2811). In total, we had eleven different mixes, with the proportion of MSI-H cells ranging from 0 to 1 (in increments of 0.1) and the remainder being MSS (Supplementary Table 1). The results of these mixing experiments were replicated 100 times, except for the pure MSS and MSI-H cases, for which all cancer cells were included. Although MSIsensor-RNA has been shown to classify single-cell RNA sequencing samples accurately [38], we checked its ability to distinguish between MSI-H and MSS samples in our datasets at the individual level. This was done by scoring individuals with MSIsensor-RNA using all available cells and again with just the cancer cells. We used the AggregateExpression function in Seurat to create the two different scenarios, and measured MSIsensor-RNA performance with ROC-AUC using the MLeval and caret packages in R [40, 41].

### 2.4 Statistical analysis

All statistical analyses were carried out in R (R version 4.1.1; https://www.R-project.org/) and all plots were created with ggplot2 [42]. Two statistical tests were performed as part of our computational pipeline. The first is a one-way ANOVA test that we used to measure ITH by comparing the difference in means between clusters of cancer cells. This was done with the aov function and was followed with Tukey’s Honestly Significant Difference test using the TukeyHSD function, both of which are from the “stats” package [43].

### 2.5 Data and code availability

All data used in this study are available either publicly through the SRA (RRID:SCR001370) and Gene Expression Omnibus (RRID:SCR005012) or through the European Genome-Phenome Archive (RRID:SCR004944) with data access requests. The associated dataset IDs and metadata for each dataset are in 1.

A distributable version of the computational pipeline, SINGLE-MSI, used in this study is available on GitHub (https://github.com/harrison-anth/single_msi). We have written the entire workflow in Snakemake to ensure reproducibility and scalability. All original results and code used in this study as well as the R script used to mix together single-cell sequencing samples are stored in another GitHub repository (https://github.com/harrison-anth/singlemsi_legacy).

## 3 Results

### 3.1 Computational pipeline distinguishes MSI-H and MSS individuals and captures ITH

To determine whether MSIsensor-RNA could distinguish between MSI-H and MSS individuals, we ran it on the aggregate expression of all cells and again on only the cancer cells for each individual. As expected, MSI-H individuals generally had higher MSI scores than MSS individuals, and MSIsensor-RNA was able to broadly distinguish between the two groups (Figure 1A,B and Supplementary Figure 1A,B). These results were seen for both aggregated expression of all cells and only cancer cells, but subsetting down to only cancer cells yielded lower MSI scores. There were also disagreements between PCR/IHC MSI status and MSIsensor-RNA score with several MSS individuals having relatively high MSI scores and several MSI-H individuals having low MSI scores (Figure 1A,B). Next, we simulated different levels of heterogeneity to determine how well our pipeline captured ITH in MSI status. For this purpose, we simulated different levels of heterogeneity, ranging progressively from pure MSS cells to pure MSI-H cells by mixing together samples from two individuals comprised of homogeneously MSI-H cancer cells and homogeneously MSS cancer cells (Supplementary Tables 1 and 2). As expected, the more homogeneous samples (MSS, mix M1, mix M9 and MSI-H in Figure 1C) had low F-statistic values while mixtures with more equal proportions of MSI-H and MSS cells (mixes M3-M5 in Figure 1C) had high F-statistic values. Increasing the proportion of MSI-H cells until mix M7 resulted in an overall reduction in the number of MSS subclones identified (Figure 1D and Supplementary Tables 1 and 2), and there was one MSI-H subclone that was consistently detected after the proportion of MSI-H cells was 0.1 (mixes M1-M9). Together, these results show that the F-statistic is sensitive to ITH, and that the number of subclones can be consistently identified across replicates, providing useful context to the heterogeneity.

**Fig. 1.**
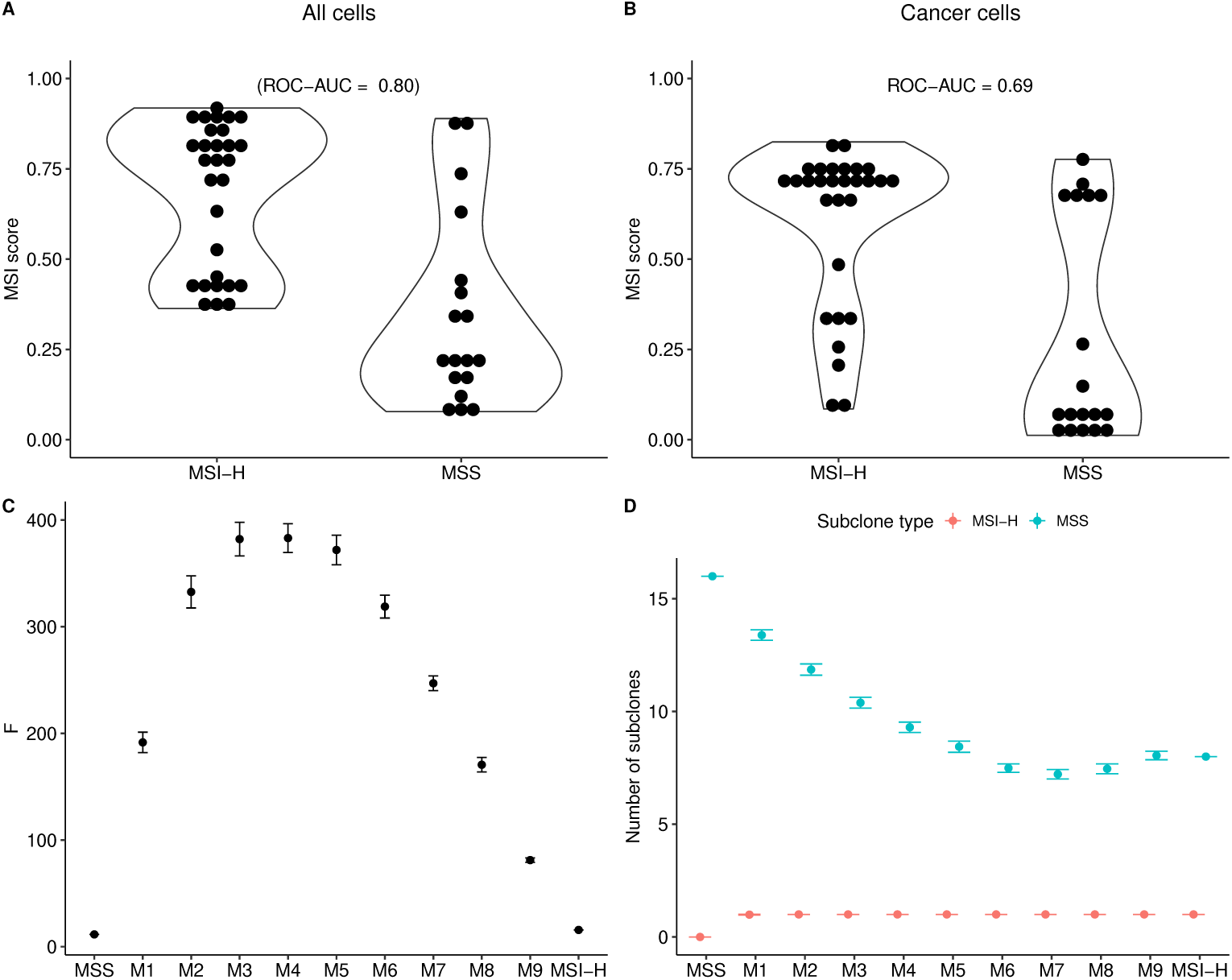
Box plots showing the distribution of (A) MSI score for individuals calculated using the aggregate expression of all cells, and (B) MSI score for individuals calculated using the aggregate expression of only cancer cells. Also shown are the mean values of (C) the F-statistic and (D) the number of subclones for the different cell mixes shown on the x-axes (with increasing proportions of MSI-H cells ranging from 0.1 in mix M1 to 0.9 in mix M9). The error bars in (C) and (D) correspond to plus/minus twice the standard error around the mean. The MSS and MSI-H samples in panels C and D are the obtained values for all cells in those samples and do not represent an average.

### 3.2 MSI-H and MSS individuals have evidence of ITH in MSI status

In order to assess heterogeneity in MSI status, we first calculated F-statistics (see Materials and Methods) based on clusters of cancer cells and identified subclones based on CNV patterns. We found that MSI-H and MSS individuals both had evidence of heterogeneity in MSI. In total, 15 of 49 individuals showed evidence of divergence in MSI status between distinct clusters of cancer cells (F *>* 25; Table 2 and Figure 2). Several individuals had very large estimates of heterogeneity based on F-statistics (75.20 to 116.10) with most of these individuals being originally deemed to be MSS, and one originally deemed to be MSI-H from a PCR or IHC test. In contrast, the lowest F-statistics (1.30 to 1.68) were found in MSI-H and MSS individuals, and the ANOVA tests were not statistically significant in either case (P *>* 0.05; Supplementary Table 3). This was also seen in most other individuals with fewer than three cancer cell clusters (Table 2 and Supplementary Table 3). In general, MSI-H and MSS individuals had similar distributions of F-statistic values but with several outliers having large F-statistic values among the MSS individuals (Figure 2A,B). Interestingly, nearly every individual in the analysis had both MSI-H and MSS subclones, and a larger proportion of MSS subclones (Figure 3 and Table 2). The exceptions were two individuals who each had a subclone proportion of 0.5 (Figure 3B), but they had very few cancer cells and too few cancer clusters to calculate an F-statistic for comparison (Supplementary Table 3). Those with the most MSI-H subclones, six and eight, were originally determined to be MSI-H, but one MSS individual also had four MSI-H subclones. Independent of MSI status, the distribution in number of clusters across individuals was relatively even, and most individuals had two or fewer samples used in the clustering process (Figure 2C,D).

**Fig. 2.**
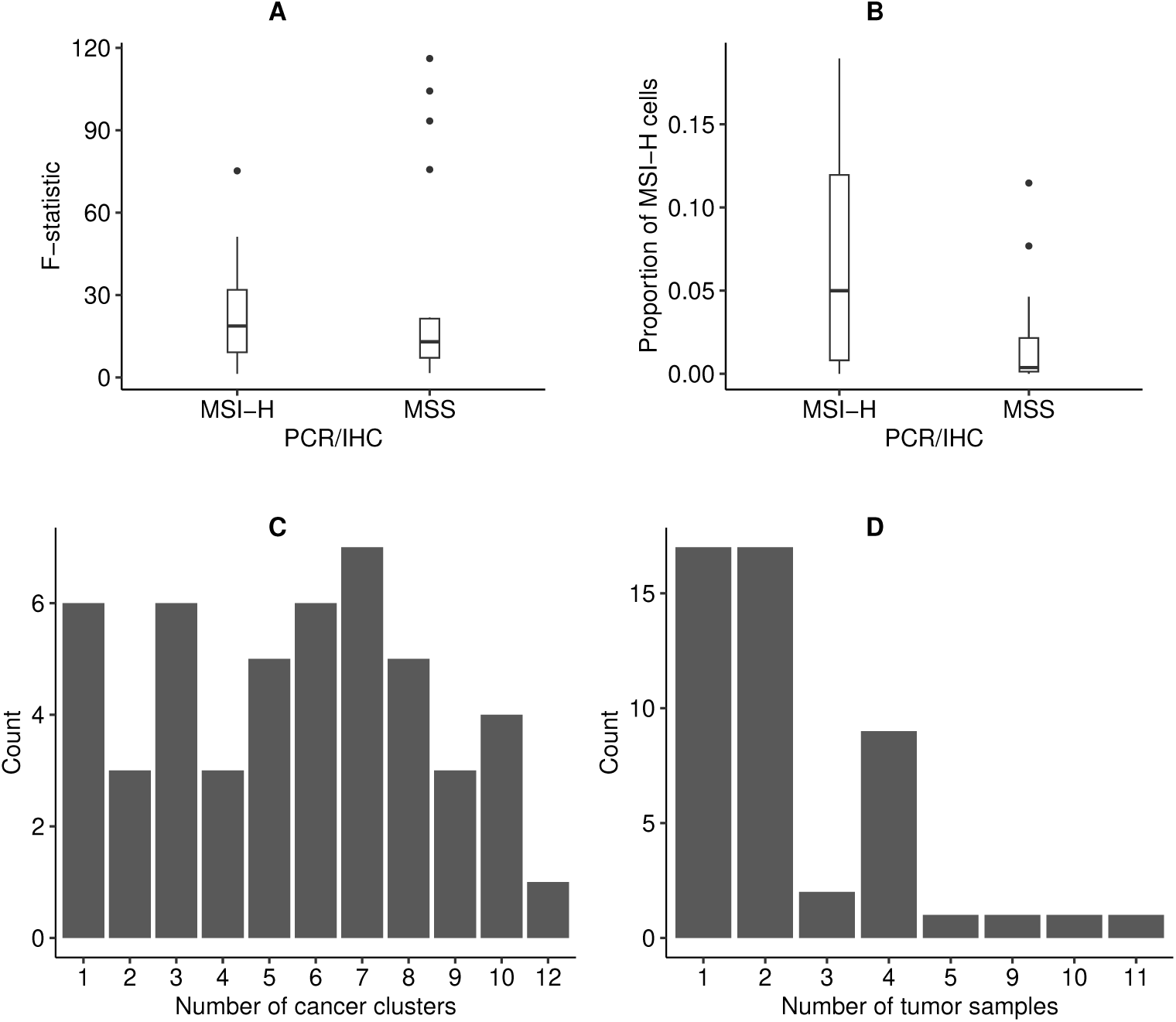
Box plots showing the distribution of (A) F-statistics grouped by PCR/IHC MSI status, (B) the proportion of MSI-H to MSS cells grouped by PCR/IHC MSI status. Also shown are histograms displaying the frequency of (C) the number of cancer cell clusters and (D) the number of tumor samples for all individuals.

**Fig. 3.**
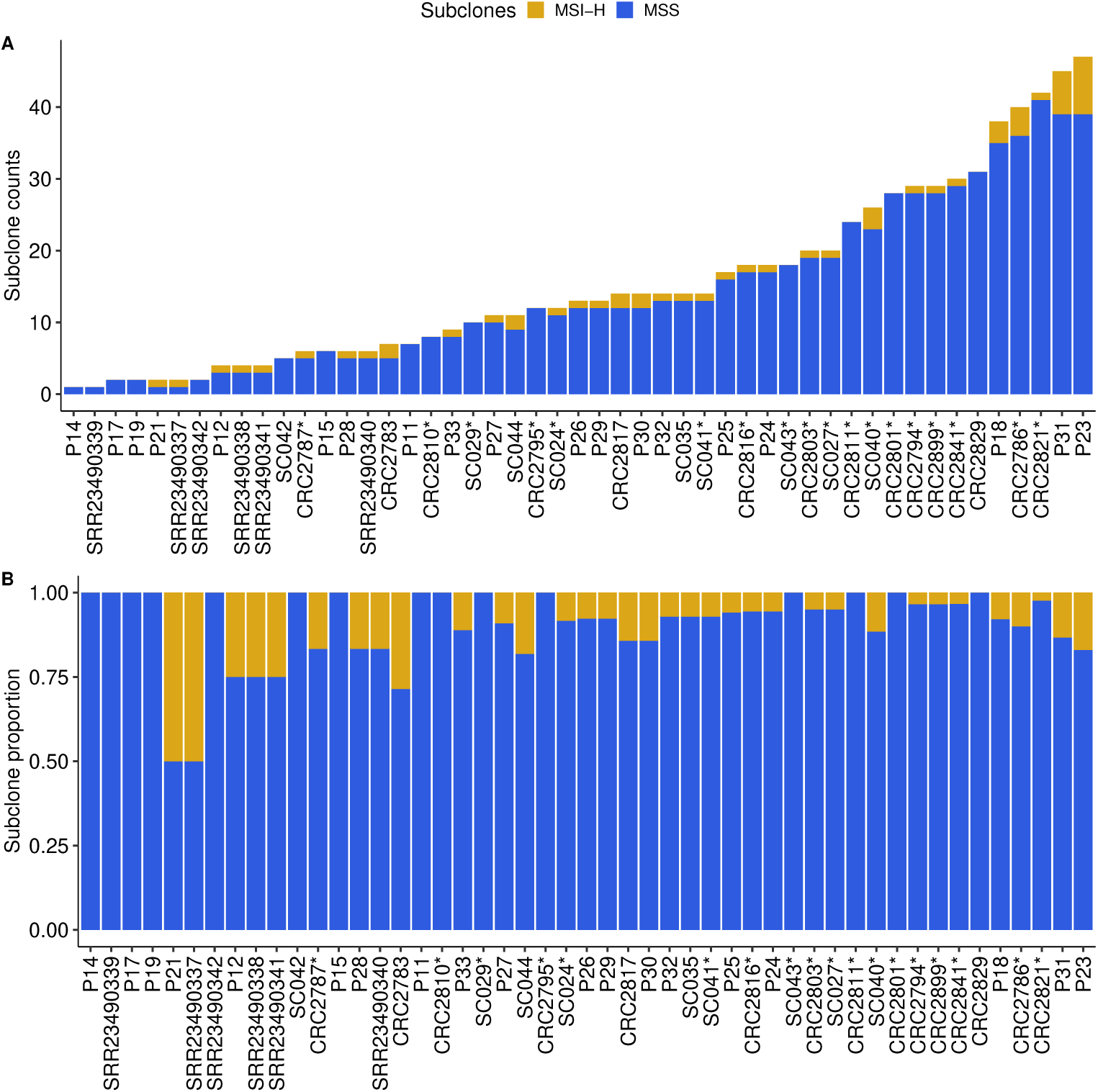
Stacked bar plots of (A) the number of subclones for each individual in the analysis and (B) the proportion of subclone types for each individual. Individuals that had a PCHR/IHC test result of MSS are indicated with an asterisk.

### 3.3 Single-cell level resolution of heterogeneity in one MSI-H and one MSS individual

We selected two individuals (P24 and CRC2786) with relatively high F-statistics and many MSI-H subclones to illustrate the heterogeneity in MSI that is evident from single cell RNA-Seq data (Figures 4,5). The MSI-H individual, P24, had good overlap in cells classified as cancer with scATOMIC (Figure 4A) and those with high MSI scores determined by MSIsensor-RNA (Figure 4B). The re-clustered cancer cells appear to cluster by MSI score; notably, clusters two and three (Figure 4C). Those larger differences in the clusters were also seen in the pseudobulk analysis of the re-clustered cancer cells (4D). The MSS individual, CRC2786, also had good overlap between the cells determined to be cancerous and those with a high MSI score; however, there was less separation of cancer and normal cells in this individual (Figure 5A,B). Similarly, in the re-clustered cancer cells, cells with higher and lower MSI scores were somewhat more intermingled than for the MSI-H individual, although clusters three and four seem to be predominantly MSS and MSI-H, respectively (Figure 5C). This result is recapitulated in the pseudobulk analysis with cluster three showing a low MSI score and cluster four a much higher one (Figure 5D).

**Fig. 4.**
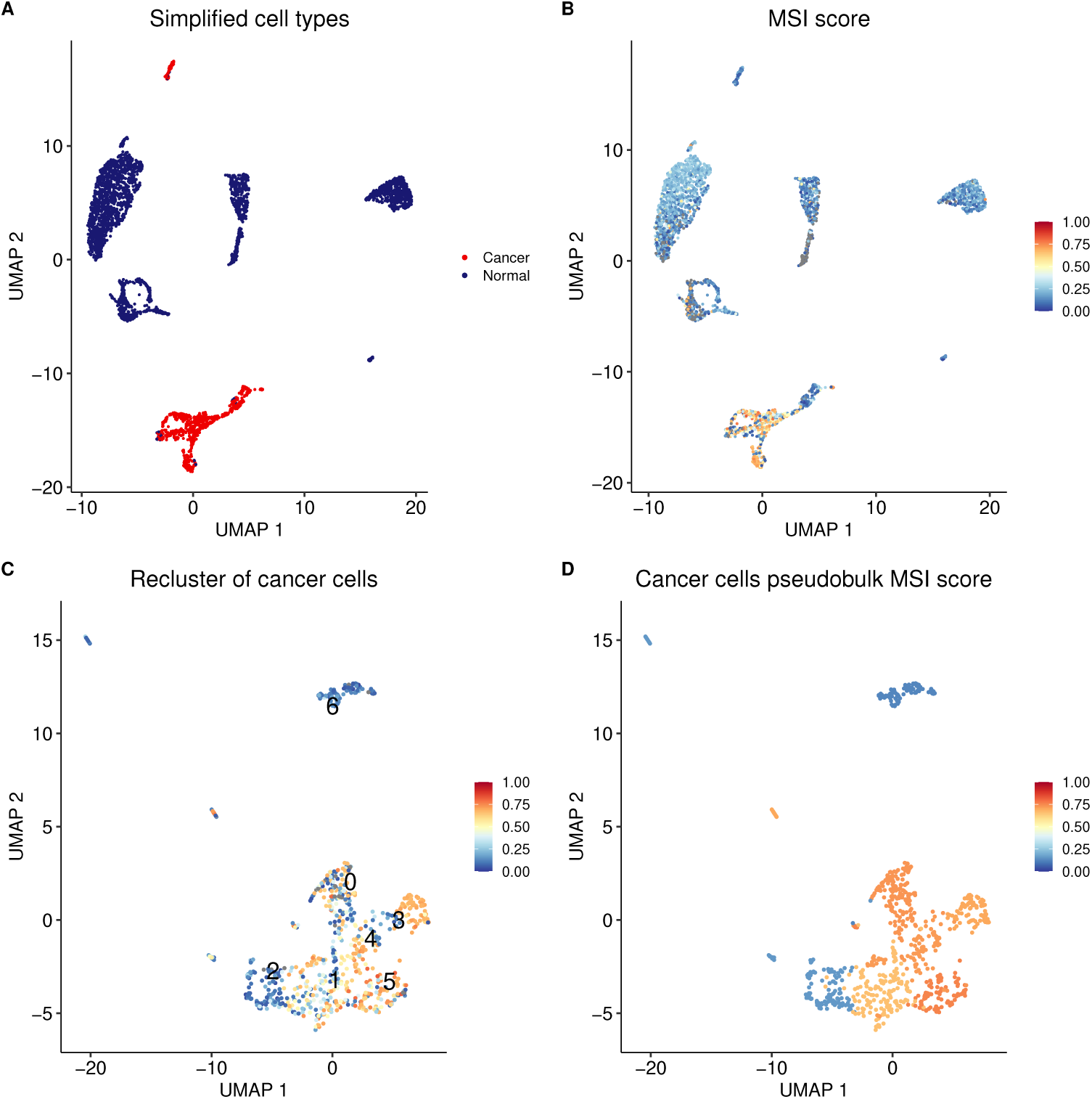
UMAP plots for MSI-H individual P24 showing (A) tumor versus normal cell classification, (B) MSI scores for each cell, (C) MSI scores for re-clustered cancer cells, and (D) MSI score for aggregated pseudobulk expression of each cancer cell cluster.

**Fig. 5.**
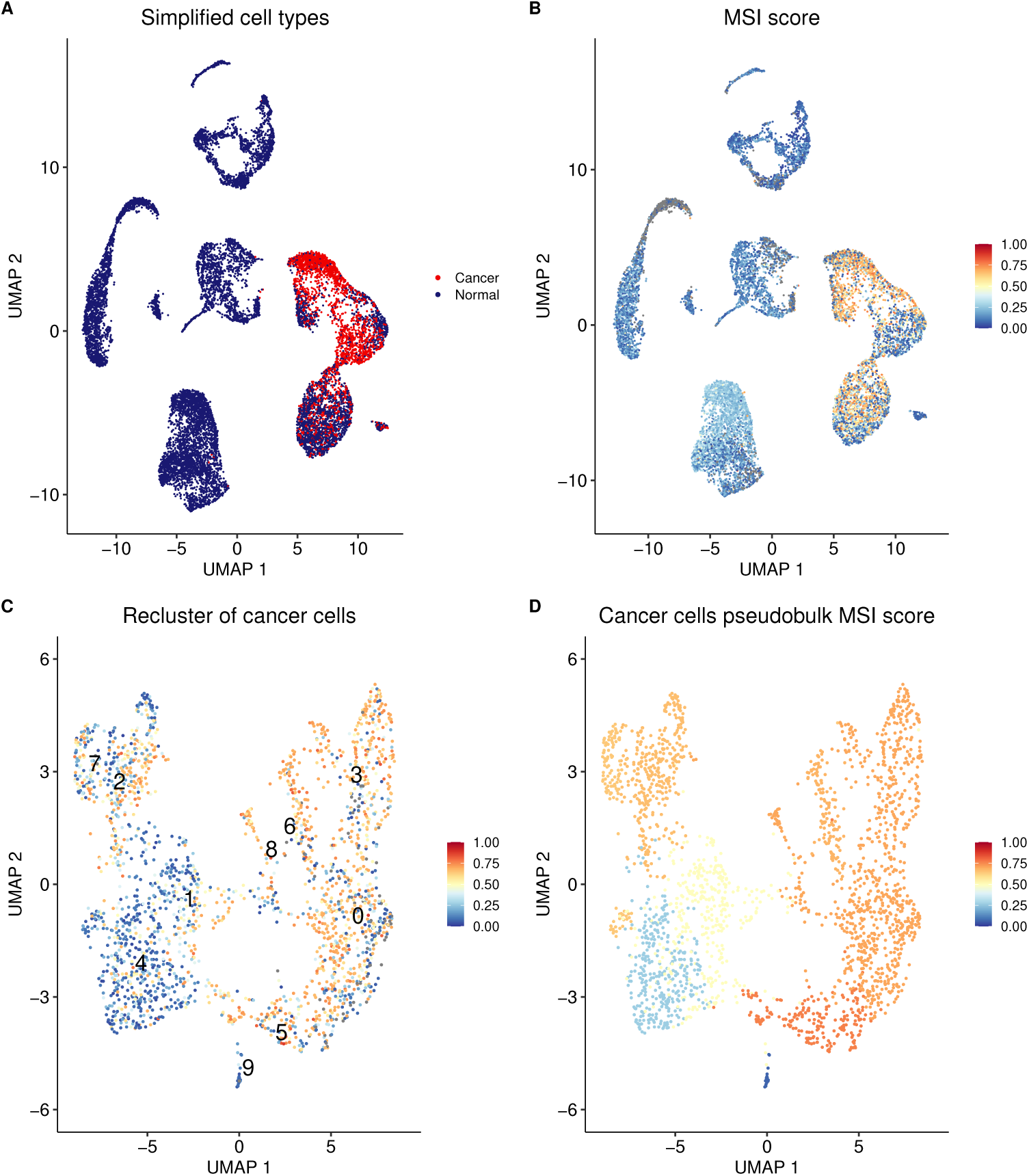
UMAP plots for MSS individual CRC2786 showing (A) tumor versus normal cell classification, (B) MSI scores for each cell, (C) MSI scores for re-clustered cancer cells, and (D) MSI score for aggregated pseudobulk expression of each cancer cell cluster.

### 3.4 Significant differences in MSI score and gene expression between clusters of cancer cells

We examined MSI ITH in CRC2786 and P24 further by assessing differences between clusters of cancer cells. We found both individuals to have many clusters with significantly different MSI scores (Figure 6), and several genes showed differences in expression between clusters and between cells identified as MSI-H and MSS (**Supplementary Figures 2**,3). Both individuals had clusters with high and low MSI scores (Figure 6A,B). These differences were found to be statistically significant (P *<* 0.05 using a Tukey HSD test; Supplementary Tables 4,5). We found that 35 cluster pairs for CRC2786 had significantly different MSI scores and 17 cluster pairs for P24 (Figure 6C,D) were significantly different. Within the clusters of cancer cells gene expression was also significantly different between clusters (Supplementary Figures 2A,3A; Supplementary Tables 6,7), and between the MSI-H and MSS cells in those clusters (Supplementary Figures 2B,3B; Supplementary Tables 8,9). The top five differentially expressed genes for each cluster of cancer cells for each individual were retained for analysis as well as the top 50 differentially expressed genes between MSI-H and MSS cells. Individual CRC2786 had three genes: *MALAT1*, *EEF1A1*, and *SH3BGRL3* in common between those differentially expressed between clusters and between cells with different MSI status; Supplementary Figure 2A,B). P24, on the other hand had one gene, *PCLAF* that was differentially expressed between clusters and between MSI-H and MSS cells. When comparing the differential expression analyses for both individuals, we found three genes: *BMX*, *LRMP*, and *SH2D6* that were differentially expressed between clusters and two genes (*TYMS*, *OXCT1* ) in common that differentiated MSI-H and MSS cells.

**Fig. 6.**
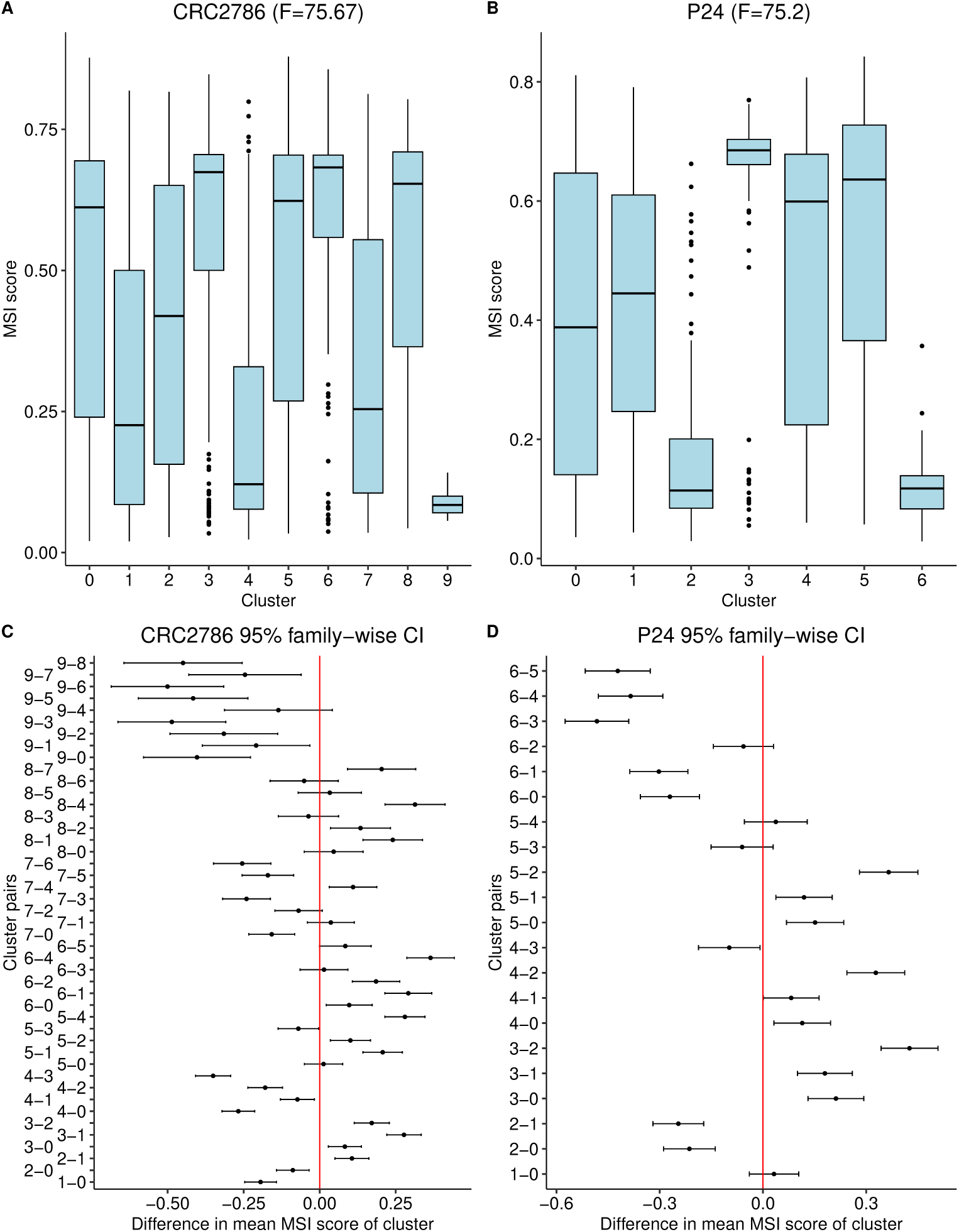
Box plots showing the distribution of MSI scores for each cluster of cancer cells in (A) individual CRC2786 and (B) individual P24. Also shown are the 95% confidence intervals for the difference in mean MSI scores between each cluster pair for (C) individual CRC2786 and (D) individual P24.

## 4 Discussion

In our study, we showed that MSI status can be heterogeneous at the single-cell level and provide a pipeline to measure that heterogeneity with the clustering of cancer cells and CNV based subclone analysis. These results contrast with the assumption that is commonly made, both in research and in clinical practice, that MSI is dichotomous. While this assumption has proven useful, enabling MSI-H to be applied as a biomarker for immune checkpoint inhibitor treatment [44], overall responder rate has been reported to be as low as 31% [29]. This could be explained, at least in part, by the heterogeneity in MSI-H individuals, which a binary classification fails to take into account. Furthermore, it is well known that single-sample tests (like the ones used to assign MSI status) are susceptible to under-sampling bias when ITH is present [3] and multi-sample, multi-regional tests may be needed to improve classification. This is supported by a recent study [45] that reported a higher accuracy in predicting immunotherapy effectiveness over traditional PCR/IHC tests by incorporating MSI cell type proportion into an MSI score. Similarly to our study, this also found that both MSI-H and MSS individuals had a mixture of MSI-H and MSS cell types in single-cell sequencing data; however, their methodology, which involved clustering cells based on gene-set enrichment of MSI-H and MSS signatures, did not identify any MSS individuals with only MSS cells. Our pipeline was able to find examples of MSS individuals comprising MSS cells only, which would make sense given that microsatellite instability is a relatively rare trait, and it would be unlikely to be present in every MSS individual in a study cohort. This is likely due to the main difference between our methods, as we test individual cells for microsatellite instability, whereas Zhao et al. labelled cells as MSI-H at the pseudo-bulk level with gene-set enrichment guided cell clustering. Our study is also different as we aimed to measure ITH and provide our pipeline in an open access format.

Our finding that nearly every MSI-H and MSS individual had MSI-H and MSS sub-clones has not yet been reported in other studies; however, two case studies that infer subclonality of dMMR status from discordant IHC test results have been reported [22, 46]. Combined with our findings, these case studies provide insights that could help explain reports of 30% or more of MSI-H cancers having primary resistance to single-agent immune checkpoint inhibitor treatment [28, 47]. A treatment regime for an MSI-H cancer would potentially miss one or more MSS subclones, leaving behind a population of cells that would not respond in the same way to immunotherapy. Although this would need to be demonstrated with a clinical experiment in which treatment results are measured longitudinally, our results provide a plausible mechanism for treatment resistance which is not currently given adequate consideration [47].

Our computational pipeline is the first to identify and quantify heterogeneity in MSI status at the single-cell level. We built the pipeline around MSIsensor-RNA and scATOMIC, two pan-cancer, machine learning based approaches. The combination of these programs may give rise to some potential issues. Naturally, as both approaches are trained on gene expression data, there will be overlap in genes used to train both classifiers and consequently overlap in cell type prediction. Yet, we found different genes to be differentially expressed between cancer cell clusters and MSI-H and MSS cells. This is likely because there is no overlap in training data between the two tools. One other caveat is the loss of microsatellite instability signal in MSI-H individuals after subsetting down to the cancer cells. Despite being necessary at the single-cell level to only label cells as MSI-H if they were also determined to be cancerous by scATOMIC, there were likely instances where MSIsensor-RNA correctly identified MSI-H cells and scATOMIC did not. Going forward, it would be beneficial for a benchmarking study to be done to determine if MSIsensor-RNA could also better identify cancer cells in MSI-H individuals. Another factor to consider is that there can be an overlap between the genes used in clustering of cells and the genes used to generate an MSI score. Whether one or more of the 100 genes used in the MSIsensor-RNA baseline are included in the 2,000 most variable genes used in clustering steps of pipeline will change from individual to individual. While not included in this study, we have checked clustering of cancer cells with and without the 100 genes used by MSIsensor-RNA and found it did not appear to affect the clustering results.

MSI is typically detected in next-generation sequencing data by comparing the distribution of insertions and deletions in microsatellites between a paired-normal and tumor sample. Because we only had access to single-cell RNA sequencing data, we chose to use MSIsensor-RNA, one of the only methods reported to accurately classify single-cell RNA sequencing samples. We found that it could broadly distinguish between the individuals deemed MSI-H and MSS with PCR/IHC tests (Figure 1A,B and **Supplementary Figure 1A,B**). However, it is worth noting that this method does not directly detect MSI with microsatellites but uses machine learning models trained on gene expression patterns from MSI-H and MSS individuals. This technique is more suited to detecting dMMR, which is traditionally measured with differences in gene expression. However, the two are inherently related as both are used as predictive biomarkers for the same immunotherapy, and MSI is hypothesized to be the down-stream result of dMMR. Based on these differences, it would be worthwhile to replicate these findings with DNA based assays (like whole-genome amplification and sequencing) which would permit the use of NGS tools that measure differences in microsatellite repeats.

## 5 Conclusions

We provide evidence of ITH in MSI status at the single-cell level and provide a pipeline that is able to identify and quantify it. Importantly, our results suggest that heterogeneity in MSI status is more common than previously reported, and it was found in both MSI-H and MSS individuals. These results could help to explain why there are reports of treatment resistance and low response rates in MSI-H cancers treated with immune checkpoint inhibitors. However, our study only analyzed data from 49 individuals that underwent single-cell RNA sequencing. Further studies are warranted to determine the frequency of heterogeneity in this biomarker at the population level and whether the presence of MSI-H and MSS subclones can have clinical impacts, including the capacity for rapid evolution of resistance to treatments for which MSI-H is used as a biomarker.

## Acknowledgements

We would like to thank the patients and researchers who made this study possible by sharing their data. This includes patients from the CRC-SG1, KUL3 and KUL5 cohorts in Joanito et al. 2022, patients involved in PICC study (NCT03926338) from Li et al. 2023, and the 6 individuals from Yunnan Cancer Hospital from Wu et al. 2023. We would also like to thank Michéal Ó Däalaigh for useful conversations on navigating single-cell cancer data and Anna Großbach for advice on graphic design. Additionally, this research was funded by Research Ireland through the Research Ireland Centre for Research Training in Genomics Data Science under Grant number 18/CRT/6214.

## Supplementary information

**Supplementary Table 1:** Summary statistics for all mixing runs reporting the average F-statistic, number of MSI-H subclones, and number of MSS subclones. The MSI-H and MSS mixes are not representative of an average as they are the results for samples used in the mixing experiment. Also reported is twice the standard error (SE) for all the summary statistics. (filename: supplementary table1.csv)

**Supplementary Table 2:** All results from mixing experiment across 100 replicates. Clusters refers to the number of cancer cell clusters and cells is the number of cancer cells. MSS, MSI-H, and total describe the number of subclones identified in the analysis. (Filename: supplementary table2.csv)

**Supplementary Table 3:** ANOVA test results for each individual. We have abbreviated degrees of freedom to Df, sum of squares to Ssq, mean sum of squares to Msq, F is the ANOVA F-statistic, and Adjusted P is the p-value associated with the test. (Filename: supplementary table3.csv)

**Supplementary Table 4:** This table contains the Tukey HSD results for individual P24. The Lower and Upper columns contain values describing the Tukey test boundaries. The Adjusted P column is the p-value associated with the test and ”Cluster pair” describes which two clusters are being compared (e.g. 1-0 is between clusters 1 and 0). (Filename: supplementary table4.csv)

**Supplementary Table 5:** This table contains the Tukey HSD results for individual CRC2786. The Lower and Upper columns contain values describing the Tukey test boundaries. The Adjusted P column is the p-value associated with the test and ”Cluster pair” describes which two clusters are being compared (e.g. 1-0 is between clusters 1 and 0). (Filename: supplementary table5.csv)

**Supplementary Table 6:** Results of the differential gene expression analysis between clusters of cancer cells for MSI-H individual, P24. P, is the p-value associated with the test. The column ”Average Log2FC” is the Log fold-change of the average expression between the cancer cell cluster (identified in the Cluster column) and all other clusters. The ”Percent1” and ”Percent2” columns describe the percentage of cells in which the Gene is detected in the cancer cell cluster or all other clusters, respectively. The ”Adjusted P” column is the p-value after Bonferroni correction. (Filename: supplementary table6.csv)

**Supplementary Table 7:** Results of the differential gene expression analysis between clusters of cancer cells for MSS individual, CRC2786. P, is the p-value associated with the test. The column ”Average Log2FC” is the Log fold-change of the average expression between the cancer cell cluster (identified in the Cluster column) and all other clusters. The ”Percent1” and ”Percent2” columns describe the percentage of cells in which the Gene is detected in the cancer cell cluster or all other clusters, respectively. The ”Adjusted P” column is the p-value after Bonferroni correction. (Filename: supplementary table7.csv)

**Supplementary Table 8:** Results of the differential gene expression analysis between MSI-H and MSS cells for MSI-H individual, P24. P, is the p-value associated with the test. The column ”Average Log2FC” is the Log fold-change of the average expression between MSI-H and MSS cells (identified in the Cluster column). The ”Percent1” and ”Percent2” columns describe the percentage of cells in which the Gene is detected, where Percent1 will describe the the cell type identified in the Cluster column. The ”Adjusted P” column is the p-value after Bonferroni correction. (Filename: supplementary table8.csv)

**Supplementary Table 9:** Results of the differential gene expression analysis between MSI-H and MSS cells for MSS individual, CRC2786. P, is the p-value associated with the test. The column ”Average Log2FC” is the Log fold-change of the average expression between MSI-H and MSS cells (identified in the Cluster column). The ”Percent1” and ”Percent2” columns describe the percentage of cells in which the Gene is detected, where Percent1 will describe the the cell type identified in the Cluster column. The ”Adjusted P” column is the p-value after Bonferroni correction. (Filename: supplementary table9.csv)

**Supplementary Figure 1:** Plots showing performance of MSIsensor-RNA based on (A) receiver operating characteristic and (B) precision-recall curves. (Filename: supplementary figure1.pdf)

**Supplementary Figure 2:** Heatmaps of differential gene expression analysis for individual P24. Panel (A) shows differential gene expression between cancer cell clusters, and panel (B) shows differential gene expression between MSI-H and MSS cells. (Filename: supplementary figure2.pdf)

**Supplementary Figure 3:** Heatmaps of differential gene expression analysis for individual CRC2786. Panel (A) shows differential gene expression between cancer cell clusters, and panel (B) shows differential gene expression between MSI-H and MSS cells. (Filename: supplementary figure3.pdf)

