## Supplementary figures and images for "Intratumoral heterogeneity in microsatellite instability at single cell resolution"

### Supplementary Figure 1

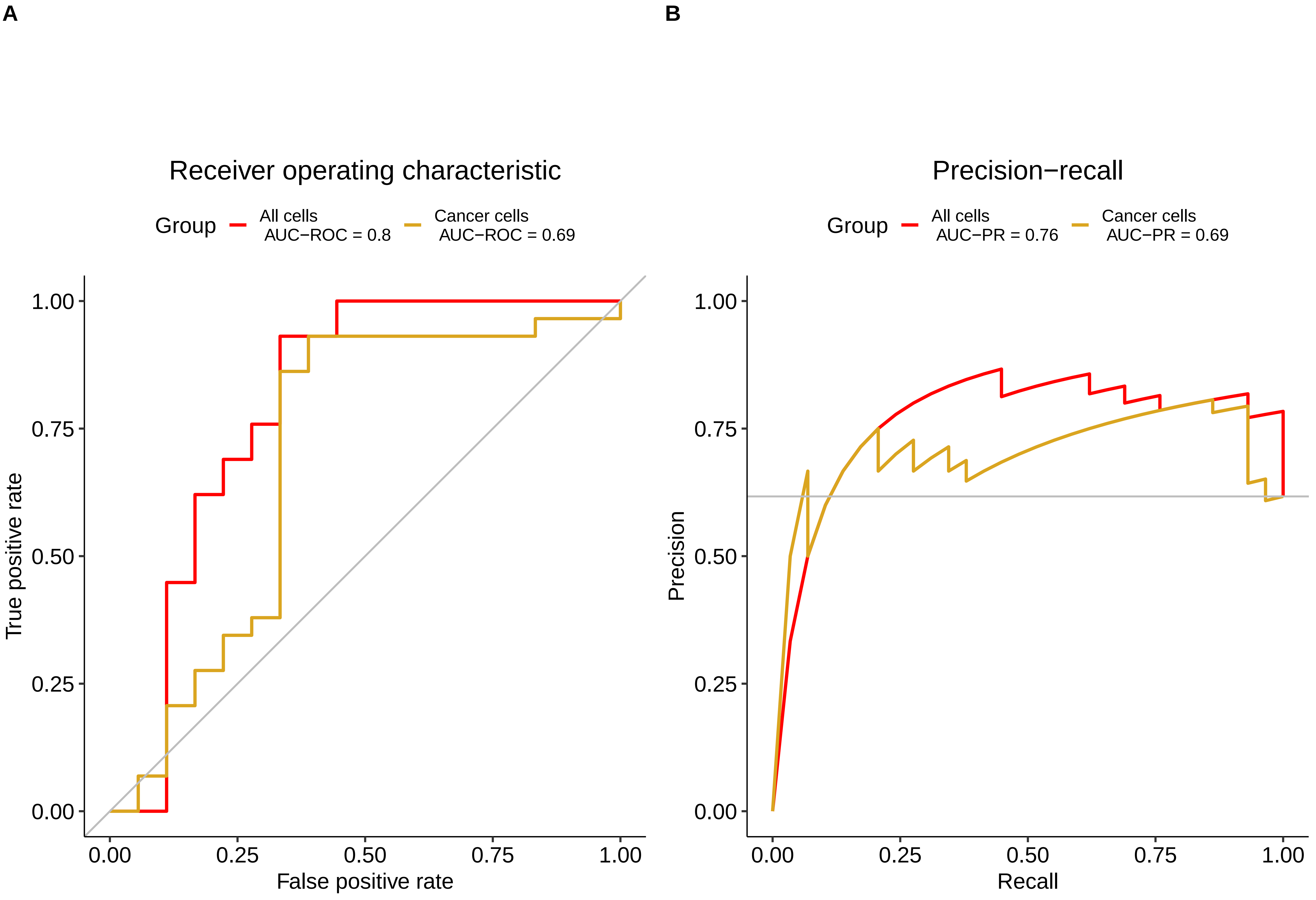

### Supplementary Figure 2

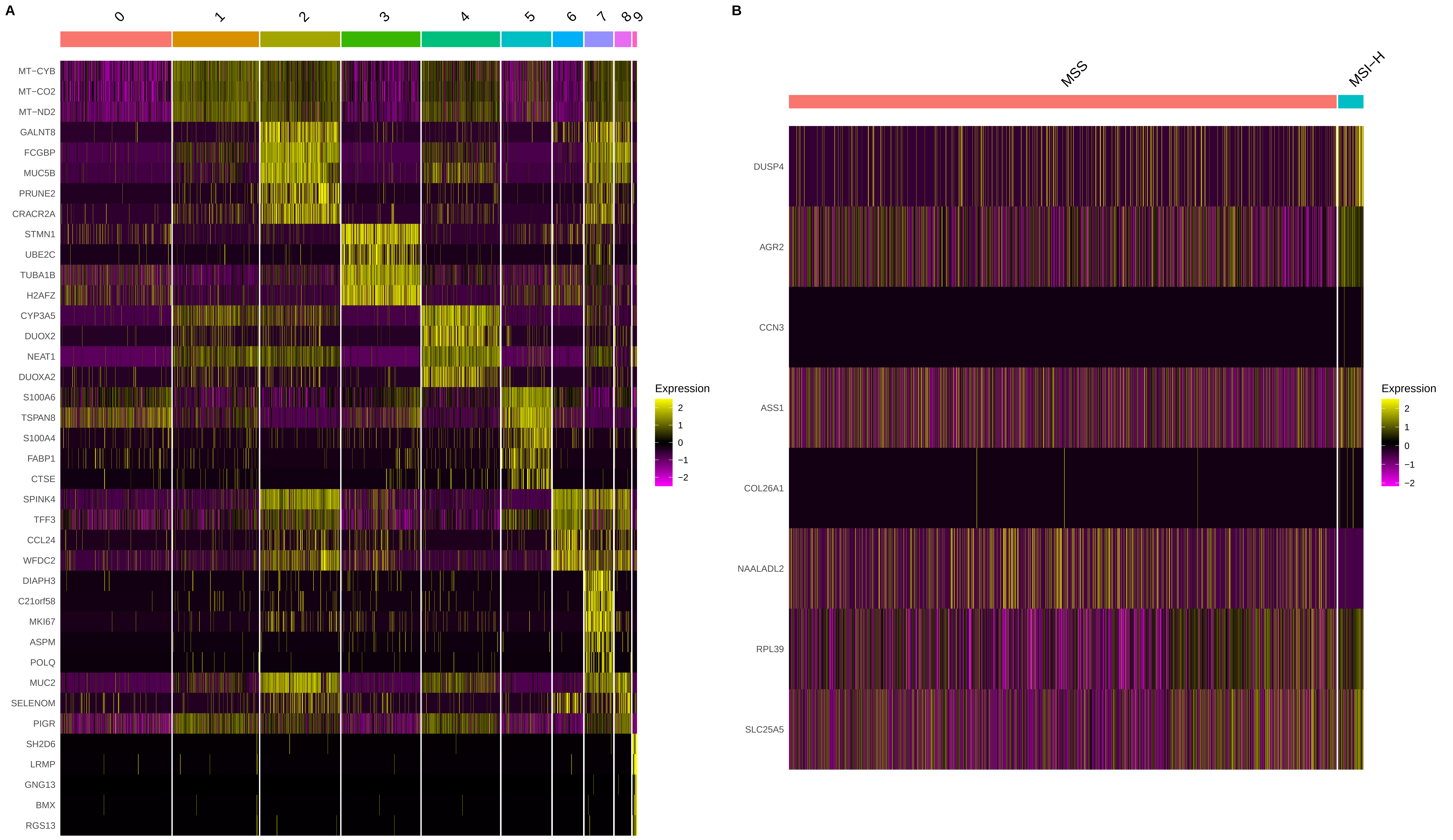

### Supplementary Figure 3

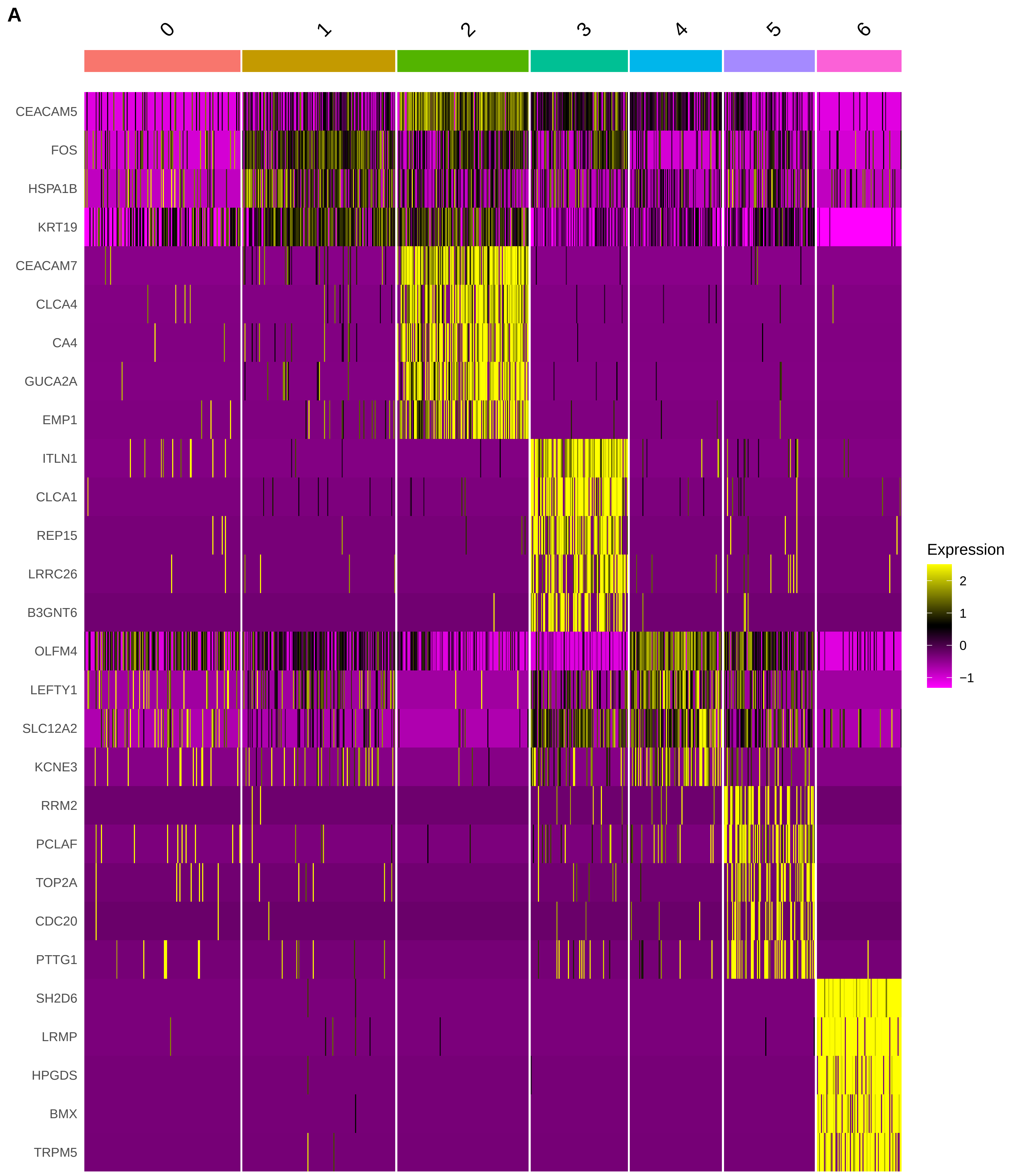

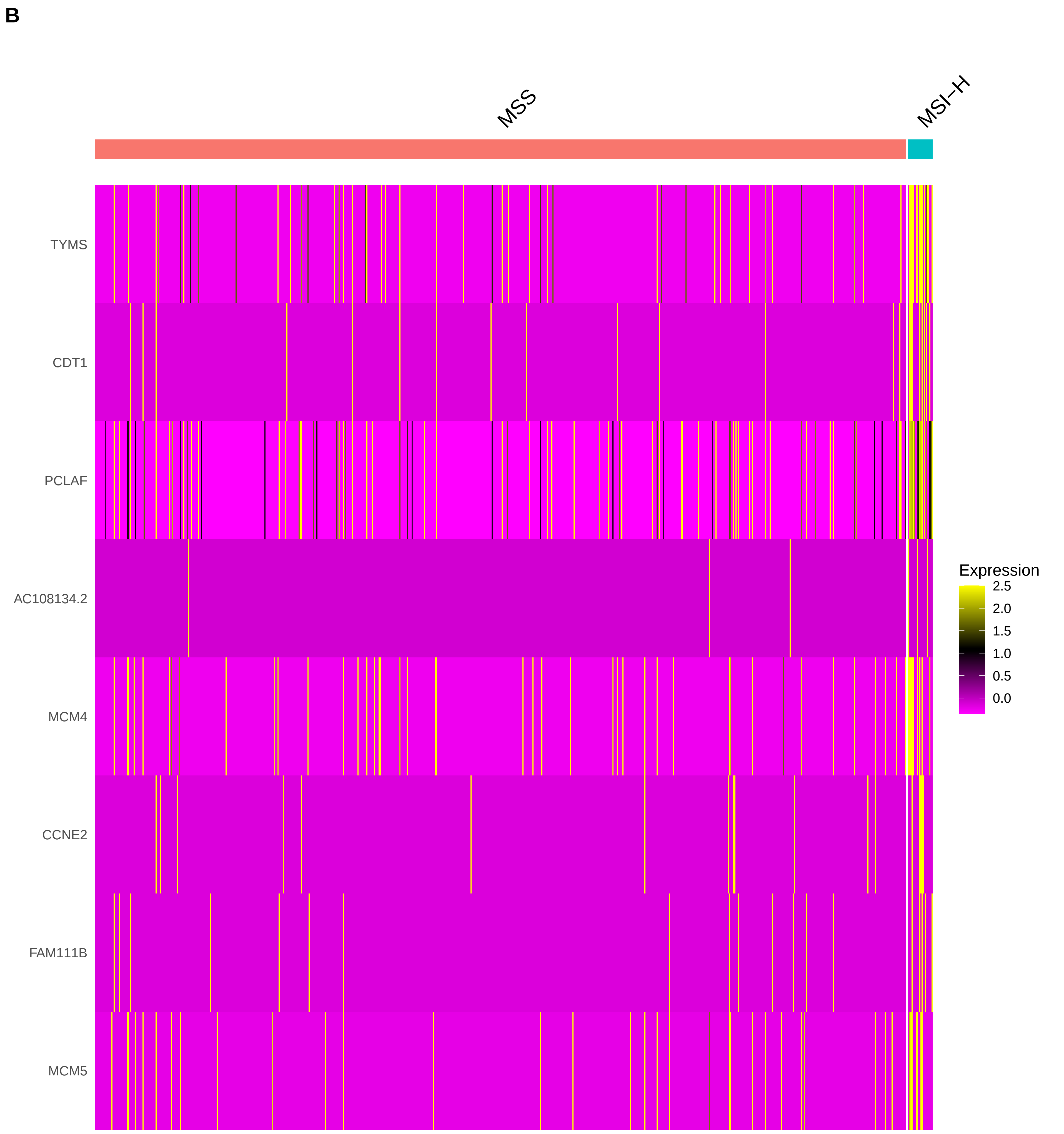
